# The Beaker Phenomenon and the Genomic Transformation of Northwest Europe

**DOI:** 10.1101/135962

**Authors:** Iñigo Olalde, Selina Brace, Morten E. Allentoft, Ian Armit, Kristian Kristiansen, Nadin Rohland, Swapan Mallick, Thomas Booth, Anna Szécsényi-Nagy, Alissa Mittnik, Eveline Altena, Mark Lipson, Iosif Lazaridis, Nick Patterson, Nasreen Broomandkhoshbacht, Yoan Diekmann, Zuzana Faltyskova, Daniel Fernandes, Matthew Ferry, Eadaoin Harney, Peter de Knijff, Megan Michel, Jonas Oppenheimer, Kristin Stewardson, Alistair Barclay, Kurt W. Alt, Azucena Avilés Fernández, Eszter Bánffy, Maria Bernabò-Brea, David Billoin, Concepción Blasco, Clive Bonsall, Laura Bonsall, Tim Allen, Lindsey Büster, Sophie Carver, Laura Castells Navarro, Oliver Edward Craig, Gordon T. Cook, Barry Cunliffe, Anthony Denaire, Kirsten Egging Dinwiddy, Natasha Dodwell, Michal Ernée, Christopher Evans, Milan Kuchařík, Joan Francès Farré, Harry Fokkens, Chris Fowler, Michiel Gazenbeek, Rafael Garrido Pena, María Haber-Uriarte, Elżbieta Haduch, Gill Hey, Nick Jowett, Timothy Knowles, Ken Massy, Saskia Pfrengle, Philippe Lefranc, Olivier Lemercier, Arnaud Lefebvre, Joaquín Lomba Maurandi, Tona Majó, Jacqueline I. McKinley, Kathleen McSweeney, Mende Balázs Gusztáv, Alessandra Modi, Gabriella Kulcsár, Viktória Kiss, András Czene, Róbert Patay, Anna Endrődi, Kitti Köhler, Tamás Hajdu, João Luís Cardoso, Corina Liesau, Michael Parker Pearson, Piotr Włodarczak, T. Douglas Price, Pilar Prieto, Pierre-Jérôme Rey, Patricia Ríos, Roberto Risch, Manuel A. Rojo Guerra, Aurore Schmitt, Joël Serralongue, Ana Maria Silva, Václav Smrčka, Luc Vergnaud, João Zilhão, David Caramelli, Thomas Higham, Volker Heyd, Alison Sheridan, Karl-Göran Sjögren, Mark G. Thomas, Philipp W. Stockhammer, Ron Pinhasi, Johannes Krause, Wolfgang Haak, Ian Barnes, Carles Lalueza-Fox, David Reich

**Author notes:** Principal investigators who contributed centrally to this study (Ian Armit, Kristian Kristiansen, Wolfgang Haak, Ian Barnes, Carles Lalueza-Fox, David Reich). To whom correspondence should be addressed: D.R., I.O.

## Abstract

Bell Beaker pottery spread across western and central Europe beginning around 2750 BCE before disappearing between 2200–1800 BCE. The mechanism of its expansion is a topic of long-standing debate, with support for both cultural diffusion and human migration. We present new genome-wide ancient DNA data from 170 Neolithic, Copper Age and Bronze Age Europeans, including 100 Beaker-associated individuals. In contrast to the Corded Ware Complex, which has previously been identified as arriving in central Europe following migration from the east, we observe limited genetic affinity between Iberian and central European Beaker Complex-associated individuals, and thus exclude migration as a significant mechanism of spread between these two regions. However, human migration did have an important role in the further dissemination of the Beaker Complex, which we document most clearly in Britain using data from 80 newly reported individuals dating to 3900–1200 BCE. British Neolithic farmers were genetically similar to contemporary populations in continental Europe and in particular to Neolithic Iberians, suggesting that a portion of the farmer ancestry in Britain came from the Mediterranean rather than the Danubian route of farming expansion. Beginning with the Beaker period, and continuing through the Bronze Age, all British individuals harboured high proportions of Steppe ancestry and were genetically closely related to Beaker-associated individuals from the Lower Rhine area. We use these observations to show that the spread of the Beaker Complex to Britain was mediated by migration from the continent that replaced >90% of Britain’s Neolithic gene pool within a few hundred years, continuing the process that brought Steppe ancestry into central and northern Europe 400 years earlier.

---

During the third millennium Before the Common Era (BCE), two new archaeological pottery styles expanded across Europe, replacing many of the more localized styles that preceded them^1^. The “Corded Ware Complex” in central, northern and eastern Europe was associated with people who derived most of their ancestry from eastern European Yamnaya steppe pastoralists^2–4^. Bell Beaker pottery is known from around 2750 cal BCE^5,6^ in Atlantic Iberia, although its exact origin is still a matter of debate^7,8^. By 2500 BCE, it is possible to distinguish in many regions the “Beaker Complex”, defined by assemblages of grave goods including stylised bell-shaped pots, distinctive copper daggers, arrowheads, stone wristguards and V-perforated buttons^9^. Regardless of the geographic region where it originated (if it did have a single origin), elements of the Beaker Complex rapidly spread throughout western Europe (and northern Africa), reaching southern and Atlantic France, Italy and central Europe^10–12^ where they overlapped geographically with the Corded Ware Complex, and from there expanding to Britain and Ireland^13,14^. A major debate has centred on whether the spread of the Beaker Complex was mediated by the movement of people, culture, or a combination of these^15–18^. Genome-wide data have revealed high proportions of Steppe ancestry in Beaker Complex-associated individuals from Germany and the Czech Republic^2–4^, consistent with their being a mixture of populations from the Steppe and the preceding farmers of Europe. However, a deeper understanding of the ancestry of people associated with the Beaker Complex requires genomic characterization of individuals across the geographic range and temporal duration of this archaeological phenomenon.

## Ancient DNA data and authenticity

To understand the genetic structure of ancient people associated with the Beaker Complex and their relationship to preceding, subsequent and contemporary peoples, we enriched ancient DNA libraries for sequences overlapping 1,233,013 single nucleotide polymorphisms (SNPs) by hybridization DNA capture^4,19^, and generated new sequence data from 170 ancient Europeans dating to ∼4700–1200 BCE (Supplementary Table 1; Supplementary Information, section 1). We also generated 62 new direct radiocarbon dates (Extended Data Table 1). We filtered out libraries with low coverage (<10,000 SNPs) or evidence of contamination (Methods) to obtain a final set of 166 individuals: 97 Beaker-associated individuals and 69 from other ancient populations (Fig. 1b; Extended Data Table 2), including 61 individuals from Neolithic and Bronze Age Britain. We combined our data with previously published ancient DNA data^2–4,20–37^ to form a genome-wide dataset of 476 ancient individuals (Supplementary Table 1). The combined dataset included Beaker-associated individuals from Iberia (n=20), southern France (n=4), northern Italy (n=1), central Europe (n=56), The Netherlands (n=9) and Britain (n=19). We further merged these data with 2,572 present-day individuals genotyped on the Affymetrix Human Origins array^22,31^ and 300 high coverage genomes sequenced as part of the Simons Genome Diversity Project^38^.

**Figure 1.**
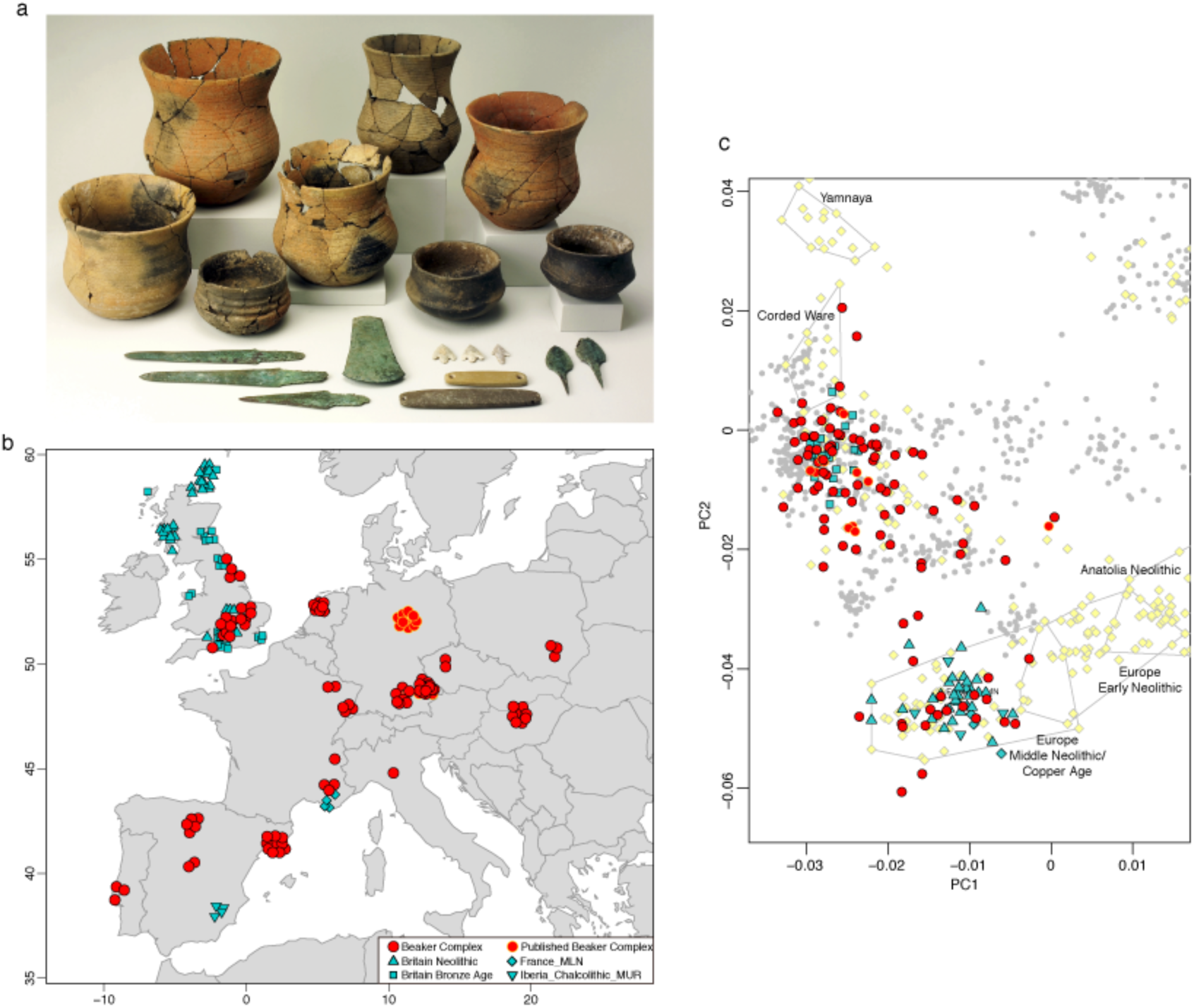
Genetic structure of individuals included in this study. **a,** Beaker Complex grave goods from La Sima III barrow^73^. Photo: Alejandro Plaza, Museo Numantino. **b**, Geographic distribution of samples with new genome-wide data, with random jitter added for clarity. **c**, Principal component analysis of 990 present-day West Eurasian individuals (grey dots), with previously published (pale yellow) and new ancient samples projected onto the first two principal components. This figure is a zoom of Extended Data Fig 1a.

## Y-chromosome analysis

We determined Y-chromosome haplogroups for the 54 male Beaker-associated individuals (Supplementary Table 3). Individuals from the Iberian Peninsula carried Y haplogroups known to be common across Europe during the earlier Neolithic period^2,4,20,26,32,39^, such as I2a (n=3) and G2 (n=1) (Supplementary Table 3). In contrast, Beaker-associated individuals outside Iberia (n=44) largely carried R1b lineages (84%), associated with the arrival of Steppe migrants in central Europe during the Late Neolithic/Early Bronze Age^2,3^. For individuals in whom we could determine the R1b subtype (n=22), we found that all but one had the derived allele for the R1b-S116/P312 polymorphism, which defines the dominant subtype in western Europe today^40^. Finding this early predominance of the R1b-S116/P312 polymorphism in ancient individuals from central and northwestern Europe suggests that people associated with the Beaker Complex may have had an important role in the dissemination of this lineage throughout most of its present-day distribution.

## Genomic insights into the spread of the Beaker Complex

Principal component analysis (PCA) revealed striking heterogeneity among individuals assigned to the Beaker Complex (Fig. 1c, Extended Data Fig. 1a). Genetic differentiation in our dataset was mainly driven by variable amounts of Steppe-related ancestry, with Beaker Complex individuals falling along the axis of variation defined by Yamnaya steppe pastoralists and Middle Neolithic/Copper Age European populations. We obtained qualitatively consistent inferences using ADMIXTURE model-based clustering^41^ (Extended Data Fig. 1b).

We grouped Beaker Complex individuals based on geographic proximity and genetic similarity (Supplementary Information, section 4), and used *qpAdm*^2^ to model their ancestry as a mixture of western European hunter-gatherers (WHG), northwestern Anatolian farmers, and Yamnaya steppe pastoralists (the first two of which contributed to earlier European farmers; Supplementary Information, section 6). We find that the great majority of Beaker Complex individuals outside of Iberia derive a large portion of their ancestry from Steppe populations (Fig. 2a), whereas in Iberia, such ancestry is absent in all sampled individuals, with the exception of two (I0461 and I0462) from the Arroyal I site in northern Spain. We detect striking differences in ancestry not only at a pan-European scale, but also within regions and even within sites. Unlike other individuals from the Upper Alsace region of France (n=2), an individual from Hégenheim resembles previous Neolithic populations and can be modelled as a mixture of Anatolian Neolithic and western hunter-gatherers without any Steppe-related ancestry. Given that the radiocarbon date of the Hégenheim individual is older (2832–2476 cal BCE (quoting 95.4% confidence intervals for this and other dates) (Supplementary Information, section 1) than other samples from the same region (2566–2133 cal BCE), the pattern could reflect temporal differentiation. At Szigetszentmiklós in Hungary, we find Beaker Complex-associated individuals with very different proportions (from 0% to 74%) of Steppe ancestry but overlapping dates. This genetic heterogeneity is consistent with early stages of mixture between previously established European farmers and migrants with Steppe ancestry. An implication is that, even at a local scale, the Beaker Complex was associated with people of diverse ancestries.

**Figure 2.**
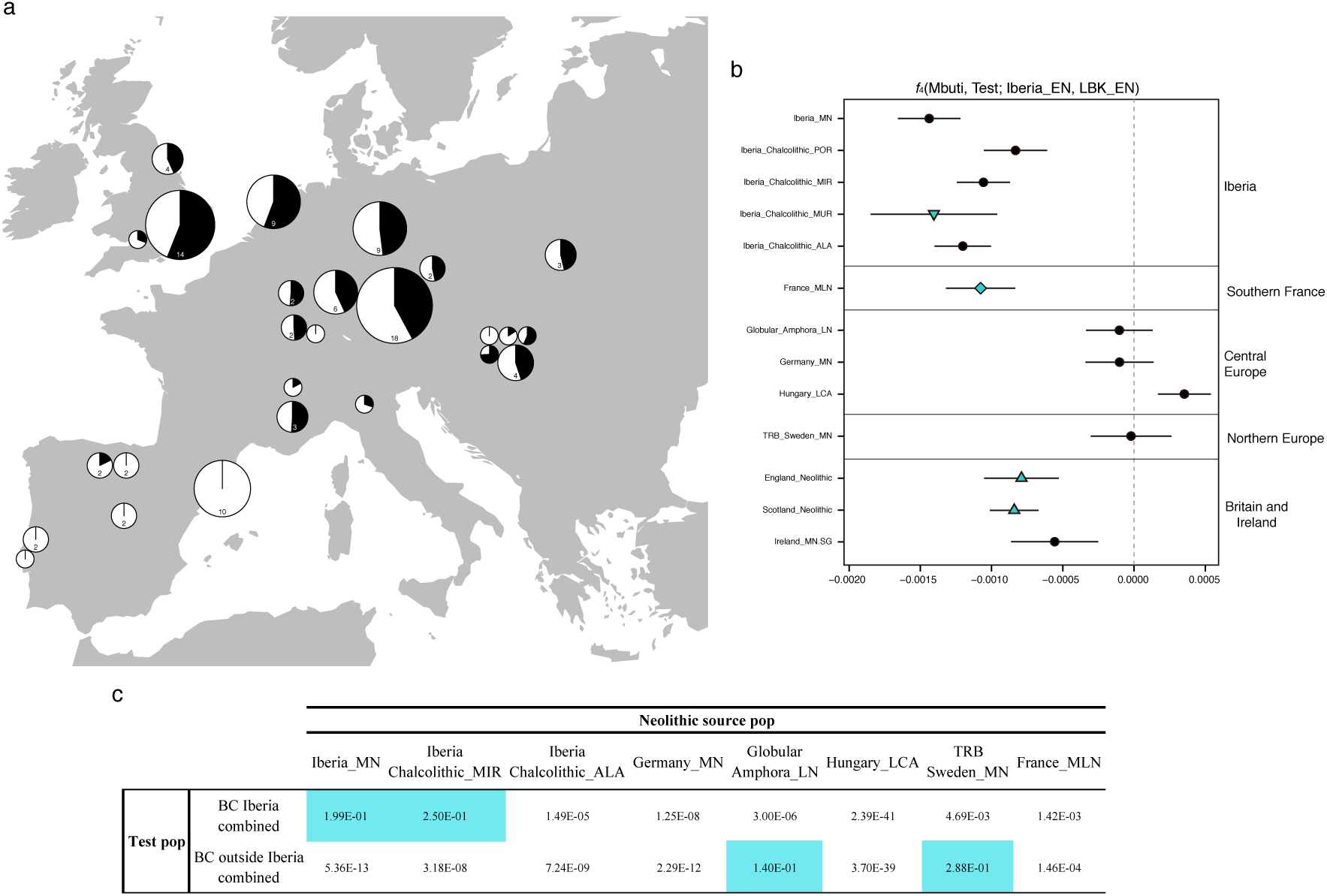
Investigating the genetic makeup of Beaker Complex individuals. **a**, Proportion of Steppe-related ancestry (shown in black) in Beaker Complex groups, computed with *qpAdm* under the model Yamnaya_Samara + Anatolia_Neolithic *+* WHG. The area of the pie is proportional to the number of individuals (shown inside the pie if more than one). See Supplementary Information, section 6 for mixture proportions and standard errors. **b**, *f-*statistics of the form *f*_4_(Mbuti, *Test*; Iberia_EN, LBK_EN) computed for European populations before the emergence of the Beaker Complex. Error bars represent ±1 standard errors. **c**, Testing different populations as a source for the Neolithic farmer ancestry component in Beaker Complex individuals. The table shows the P-values (highlighted if <0.05) for the model: Yamnaya_Samara + Neolithic farmer population. BC, Beaker complex.

While the Yamnaya-related ancestry in Beaker Complex associated individuals had an origin in the Steppe^2,3^, the other ancestry component (from European Neolithic farmers) could potentially be derived from several parts of Europe, as genetically closely related populations were widely distributed across the continent during the Neolithic and Copper Age periods^2,4,22,25,26,28,32^. To obtain insight into the origin of the Neolithic-related ancestry in Beaker Complex-associated individuals, we began by looking for regional patterns of genetic differentiation within Europe during the Neolithic and Copper Age periods. To study genetic affinity to different Early Neolithic (EN) populations, we computed *f*_4_-statistics of the form *f*_4_(*Outgroup*, *Test*; *Iberia_EN*, *LBK_EN*) for Neolithic and Copper Age test populations predating the emergence of the Beaker Complex. As previously described^2^, there is genetic affinity to Iberian Early Neolithic farmers in Iberian Middle Neolithic/Copper Age populations, but not in central and northern European Neolithic populations (Fig. 2b), which could be explained by differential affinities to hunter-gatherer individuals from different regions^42^ (Extended Data Fig. 2). A new finding that emerges from our analysis is that Neolithic individuals from southern France and Britain also show a greater affinity to Iberian Early Neolithic farmers than to central European Early Neolithic farmers (Fig. 2b), similar to previous results obtained in a Neolithic farmer genome from Ireland^28^. By modelling Neolithic populations and WHG in an admixture graph framework, we replicate these results and further show that they are not driven by different proportions of hunter-gatherer admixture (Extended Data Fig. 3; Supplementary Information, section 5). Our results suggest that a portion of the ancestry of the Neolithic farmers of Britain was derived from migrants who spread along the Atlantic coast. Megalithic tombs document substantial interaction along the Atlantic façade of Europe, and our results are consistent with such interactions reflecting movements of people. More data from southern Britain (where our sampling is sparse) and nearby regions in continental Europe will be needed to fully understand the complex interactions between Britain and the continent in the Neolithic^43^.

**Figure 3.**
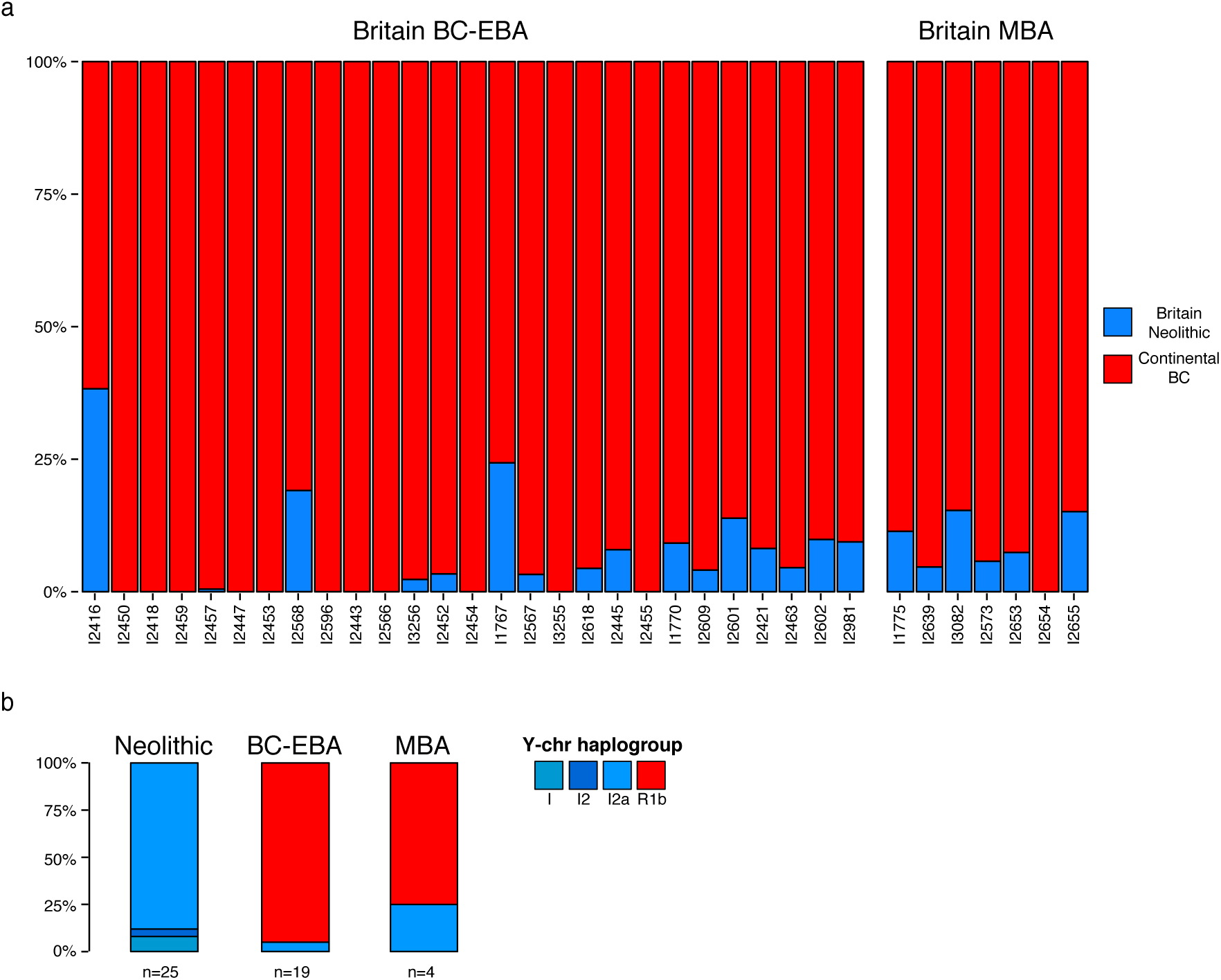
Population transformation in Britain associated with the arrival of the Beaker Complex. **a**, Modelling Beaker Complex and Bronze Age individuals from Britain as a mixture of continental Beaker Complex (red, represented by Beaker Complex samples from Oostwoud) and Britain_Neolithic (blue). Individuals are ordered chronologically (oldest on the left) and included in the plot if represented by more than 100,000 SNPs. See Supplementary Information, section 6 for mixture proportions and standard errors**. b**, Y-chromosome haplogroup distribution in males from Britain. EBA, Early Bronze Age; MBA, Middle Bronze Age. BC, Beaker complex.

The distinctive genetic signatures of pre-Beaker Complex populations in Iberia compared to central Europe allow us to test formally for the origin of the Neolithic farmer-related ancestry in Beaker Complex individuals in our dataset (Supplementary Information, section 6). We grouped individuals from Iberia (n=19) and from outside Iberia (n=84) to increase power, and evaluated the fit of different Neolithic/Copper Age groups with *qpAdm* under the model: Yamnaya + Neolithic/Copper Age. For Beaker Complex individuals from Iberia, the best fit was obtained when Middle Neolithic and Copper Age populations from the same region were used as a source for their Neolithic farmer-related ancestry, and we could exclude central and northern European populations (P < 4.69E-03) (Fig. 2c). Conversely, the Neolithic farmer-related ancestry in Beaker Complex individuals outside Iberia was most closely related to central and northern European Neolithic populations with relatively high hunter-gatherer admixture (e.g. *Globular_Amphora_LN*, P = 0.14; TRB_Sweden_MN, P = 0.29), and we could significantly exclude Iberian sources (P < 3.18E-08) (Fig. 2c). These results support largely different origins for Beaker Complex individuals, with no discernible Iberia-related ancestry outside Iberia.

## Nearly complete turnover of ancestry in Britain

British Beaker Complex individuals (n=19) show strong similarities to the central European Beaker Complex both in genetic profile (Extended Data Fig. 1) and in material culture: the great majority of individuals from both regions are associated with “All Over Corded” Beaker pottery. The presence of large amounts of Steppe-related ancestry in the British Beaker Complex (Fig. 2a) contrasts sharply with Neolithic individuals from Britain (n=35), who have no evidence of Steppe genetic affinities and cluster instead with Middle Neolithic and Copper-Age populations from mainland Europe (Extended Data Fig. 1). Thus, the arrival of Steppe ancestry in Britain was mediated by a migration that began with the Beaker Complex. A previous study showed that Steppe ancestry arrived in Ireland by the Bronze Age^28^, and here we show that – at least in Britain – it arrived by the Copper Age / Beaker period.

Among the different continental Beaker Complex groups analysed in our dataset, individuals from Oostwoud (Province of Noord-Holland, The Netherlands) are the most closely related to the great majority of the Beaker Complex individuals from southern Britain (n=14). They had almost identical Steppe ancestry proportions (Fig. 2a), the highest shared genetic drift (Extended Data Fig. 4b) and were symmetrically related to other ancient populations using *f*_4_- statistics (Extended Data Fig. 4a), showing that they are consistent with being derived from the same ancestral population without additional mixture into either group. We next investigated the magnitude of population replacement in Britain with *qpAdm*^2^ by modelling Beaker Complex and Bronze Age individuals as a mixture of continental Beaker Complex (using the Oostwoud individuals as a surrogate) and the British Neolithic population (Supplementary Information, section 6). Fig. 3a shows the results of this analysis, ordering individuals by date and showing excess Neolithic ancestry compared to continental Beaker Complex as a baseline. For the earliest individuals (between ∼2400–2000 BCE), the Neolithic ancestry excess is highly variable, consistent with migrant communities who were just beginning to mix with the previously established Neolithic population of Britain. During the subsequent Bronze Age we observe less variation among individuals and a modest increase in Neolithic-related ancestry (Fig. 3a), which could represent admixture with persisting populations with high levels of Neolithic-related ancestry (or alternatively incoming continental populations with higher proportions of Neolithic-related ancestry). In either case, our results imply a minimum of 93±2% local population turnover by the Middle Bronze Age (Supplementary Information, section 6). Specifically, for individuals from Britain around 2000 BCE, at least this fraction of their DNA derives from ancestors who at 2500 BCE lived in continental Europe. An independent line of evidence for population turnover comes from Y-chromosome haplogroup composition: while R1b haplogroups were completely absent in the Neolithic samples (n=25), they represent 95% and 75% of the Y-chromosomes in Beaker Complex-Early Bronze Age and Middle Bronze Age males in Britain, respectively (Fig. 3b; Supplementary Table 3).

Our genetic time transect in Britain also allowed us to track the frequencies of alleles with known phenotypic effects. Derived alleles at rs12913832 (SLC45A2) and rs16891982 (HERC2/OCA2), which contribute to reduced skin and eye pigmentation in Europeans, dramatically increased in frequency during the Beaker and Bronze Age periods (Extended Data Fig. 5). Thus, the arrival of migrants associated with the Beaker Complex significantly altered the pigmentation phenotypes of British populations. However, the lactase persistence allele at SNP rs4988235 remained at very low frequencies in our dataset both in Britain and continental Europe, showing that the major increase in its frequency in Britain, as in mainland Europe, occurred in the last 3,500 years^3,4,39,44^.

## Discussion

The term “Bell Beaker” was introduced by late 19^th^-century and early 20^th^-century archaeologists to refer to the distinctive pottery style found across western and central Europe at the end of the Neolithic, initially hypothesized to have been spread by a genetically homogeneous group of people. This idea of a “Beaker Folk” became unpopular after the 1960s as scepticism about the role of migration in mediating change in archaeological cultures grew^45^, although J.G.D. Clark speculated that the Beaker Complex expansion into Britain was an exception^46^, a prediction that has now been borne out by ancient genomic data.

Our results clearly prove that the expansion of the Beaker Complex cannot be described by a simple one-to-one mapping of an archaeologically defined material culture to a genetically homogeneous population. This stands in contrast to other archaeological complexes analysed to date, notably the *Linearbandkeramik* first farmers of central Europe^2^, the Yamnaya of the Pontic-Caspian Steppe^2,3^, and to some extent the Corded Ware Complex of central and eastern Europe^2,3^. Instead, or results support a model in which both cultural transmission and human migration played important roles, with the relative balance of these two processes depending on the region. In Iberia, the majority of Beaker Complex-associated individuals lacked Steppe affinities and were genetically most similar to preceding Iberian populations. In central Europe, Steppe ancestry was widespread and we can exclude a substantial contribution from Iberian Beaker Complex-associated individuals, contradicting initial suggestions of gene flow between these groups based on analysis of mtDNA^47^ and dental morphology^48^. Small-scale contacts remain plausible, however, as we observe small proportions of Steppe ancestry in two individuals from northern Spain.

Although cultural transmission seems to have been the main mechanism for the diffusion of the Beaker Complex between Iberia and central Europe, other parts of the Beaker Complex expansion were driven to a substantial extent by migration, with Beaker-associated burials in southern France, northern Italy, and Britain, representing the earliest occurrence of Steppe-related ancestry so far known in all three regions. This genomic transformation is clearest in Britain due to our dense genetic time transect. The earliest Beaker pots found in Britain show influences from both the lower Rhine region and the Atlantic façade of western Europe^49^.

However, such dual influence is not mirrored in the genetic data, as the British Beaker Complex individuals were genetically most similar to lower Rhine individuals from the Netherlands. The arrival of the Beaker Complex precipitated a profound demographic transformation in Britain, exemplified by the absence of individuals in our dataset without large amounts of Steppe-related ancestry after 2400 BCE. It is possible that the uneven geographic distribution of our samples, coupled with different burial practises between local and incoming populations (cremation versus burial) during the early stages of interaction could result in a sampling bias against local individuals. However, the signal observed during the Beaker period persisted through the later Bronze Age, without any evidence of genetically Neolithic-like individuals among the 27 Bronze Age individuals we newly report, who traced more than 90% of their ancestry to individuals of the central European Beaker Complex. Thus, the genetic evidence points to a substantial amount of migration into Britain from the European mainland beginning around 2400 BCE. These results are notable in light of strontium and oxygen isotope analyses of British skeletons from the Beaker and Bronze Age periods^50^, which have provided no evidence of substantial mobility over individuals’ lifetimes from locations with cooler climates or from places with geologies atypical of Britain. However, the isotope data are only sensitive to first-generation migrants, and do not rule out movements from regions such as the lower Rhine, which is consistent with the genetic data, or from other geologically similar regions for which DNA sampling is still sparse. Further sampling of regions on the European continent may reveal additional candidate sources.

By analysing DNA data from ancient individuals we have been able to provide important constraints on the processes underlying cultural and social changes in Europe during the third millennium BCE. Our results raise new questions and motivate further archaeological research to identify the changes in social organization, technology, subsistence, climate, population sizes^51^ or pathogen exposure^52,53^ that could have precipitated the demographic changes uncovered in this study.

## Methods

### Ancient DNA analysis

We screened skeletal samples for DNA preservation in dedicated clean rooms. We extracted DNA^54–56^ and prepared barcoded next generation sequencing libraries, the majority of which were treated with uracil-DNA glycosylase to greatly reduce the damage (except at the terminal nucleotide) that is characteristic of ancient DNA^57,58^ (Supplementary Information, section 2). We initially enriched libraries for sequences overlapping the mitochondrial genome^59^ and ∼3000 nuclear SNPs using synthesized baits (CustomArray Inc.) that we PCR amplified. We sequenced the enriched material on an Illumina NextSeq instrument with 2x76 cycles, and 2x7 cycles to read out the two indices^60^. We merged read pairs with the expected barcodes that overlapped by at least 15 base pairs, mapped the merged sequences to hg19 and to the reconstructed mitochondrial DNA consensus sequence^61^ using the *samse* command in bwa (v0.6.1)^62^, and removed duplicated sequences. We evaluated DNA authenticity by estimating the rate of mismatching to the consensus mitochondrial sequence^63^, and also requiring that the rate of damage at the terminal nucleotide was at least 3% for UDG-treated libraries^63^ and 10% for non-UDG-treated libraries^64^.

For libraries that were promising after screening, we enriched in two consecutive rounds for sequences overlapping 1,233,013 SNPs (‘1240k SNP capture’)^2,19^ and sequenced 2x76 cycles and 2x7cycles on an Illumina NextSeq500 instrument. We processed the data bioinformatically as for the mitochondrial capture data, this time mapping only to the human reference genome *hg19* and merging the data from different libraries of the same individual. We further evaluated authenticity by studying the ratio of X-to-Y chromosome reads and estimating X-chromosome contamination in males based on the rate of heterozygosity^65^. Samples with evidence of contamination were either filtered out or restricted to sequences with terminal cytosine deamination to remove sequences that could have derived from modern contaminants. Finally, we filtered out from our analysis dataset samples with fewer than 10,000 targeted SNPs covered at least once and samples that were first-degree relatives of others in the dataset (keeping the sample with the larger number of covered SNPs) (Supplementary Table 1).

### Mitochondrial haplogroup determination

We used the mitochondrial capture bam files to determine the mitochondrial haplogroup of each sample with new data, restricting to reads with MAPQ≥30 and base quality ≥30. First, we constructed a consensus sequence with samtools and bcftools^66^, using a majority rule and requiring a minimum coverage of 2. We called haplogroups with HaploGrep2^67^ based on phylotree^68^ (mtDNA tree Build 17 (18 Feb 2016)). Mutational differences compared to the rCRS and corresponding haplogroups can be viewed in Supplementary Table 2.

### Y-chromosome analysis

We determined Y-chromosome haplogroups for both new and published samples (Supplementary Information, section 3). We made use of the sequences mapping to 1240k Y-chromosome targets, restricting to sequences with mapping quality ≥30 and bases with quality ≥30. We called haplogroups by determining the most derived mutation for each sample, using the nomenclature of the International Society of Genetic Genealogy (http://www.isogg.org) version 11.110 (21 April 2016). Haplogroups and their supporting derived mutations can be viewed in Supplementary Table 3.

### Merging newly generated data with published data

We assembled two datasets for population genetics analyses:

- *HO* includes 2,572 present-day individuals from worldwide populations genotyped on the Human Origins Array^22,31,69^ and 470 ancient individuals. The ancient set includes 103 Beaker Complex individuals (87 newly reported, 5 with shotgun data^3^ for which we generated 1240k capture data and 11 previously published^3,4^), 68 newly reported individuals from relevant ancient populations and 298 previously published^2–4,20–37^ individuals (Supplementary Table 1). We kept 591,642 autosomal SNPs after intersecting autosomal SNPs in the 1240k capture with the analysis set of 594,924 SNPs from Lazaridis et al.^22^.

-*HOIll* includes the same set of ancient samples and 300 present-day individuals from 142 populations sequenced to high coverage as part of the Simons Genome Diversity Project^38^. For this dataset, 1,054,671 autosomal SNPs were used, excluding SNPs of the 1240k array located on sex chromosomes or with known functional effects.

For both datasets, ancient individuals were merged by randomly sampling one read at each SNP position, discarding the first and the last two nucleotides of each read.

### Principal component analysis

We carried out principal component analysis (PCA) on the *HO* dataset using the *smartpca* program in EIGENSOFT^70^. We computed principal components on 990 present-day West Eurasians and projected ancient individuals using lsqproject: YES and shrinkmode: YES.

### ADMIXTURE analysis

We performed model-based clustering analysis using ADMIXTURE^41^ on the *HO* reference dataset, including 2,572 present-day individuals from worldwide populations and the ancient individuals. First, we carried out LD-pruning on the dataset using PLINK^71^ with the flag -- indep-pairwise 200 25 0.4, keeping 306,393 SNPs. We ran ADMIXTURE with the cross validation (--cv) flag specifying from K=2 to K=20 clusters, with 20 replicates for each value of K and keeping for each value of K the replicate with highest log likelihood. In Extended Data Fig. 1b we show the cluster assignments at K=8 of newly reported individuals and other relevant ancient samples for comparison. This value of K was the lowest for which components of Caucasus hunter-gatherers (CHG) and European hunter-gatherers were maximized.

### *f*-statistics

We computed *f*-statistics on the *HOIll* dataset using ADMIXTOOLS^69^ with default parameters (Supplementary Information, section 4). We used *qpDstat* with f4mode:Yes for *f*_4_-statistics and *qp3Pop* for outgroup *f*3-statistics. We computed standard errors using a weighted block jackknife^72^ over 5 Mb blocks.

### Inference of mixture proportions

We estimated ancestry proportions on the *HOIll* dataset using *qpAdm*^2^ and a basic set of 9 *Outgroups*: Mota, Ust_Ishim, MA1, Villabruna, Mbuti, Papuan, Onge, Han, Karitiana. For some analyses (Supplementary Information, section 6) we added additional outgroups to this basic set.

### Allele frequency estimation from read counts

We used allele counts at each SNP to perform maximum likelihood estimation of allele frequencies in ancient populations as in ref.^4^. In Extended Data Fig. 5, we show derived allele frequency estimates at three SNPs of functional importance for different ancient populations.

## Data availability

All 1240k and mitochondrial capture sequencing data is available from the European Nucleotide Archive, accession number XXXXXXXX [to be made available on publication].

Pseudo haploid genotype data is available from the Reich Lab website at [to be made available on publication].

## Acknowledgements

We thank D. Anthony, J. Koch, I. Mathieson and C. Renfrew for comments and critiques. We thank A. C. Sousa for providing geographical information on a Portuguese sample and A. Martín and L. Loe for help in contacting archaeologists. We thank M. Giesen for assisting with samples selection. We thank the Museo Arqueológico Regional de la Comunidad de Madrid for kindly allowing access to samples from Camino de las Yeseras. We thank the Hunterian Museum, University of Glasgow, for allowing access to samples from sites in Scotland, and particularly to Dr. S.-A. Coupar for help in accessing material. We thank the Museu Municipal de Torres Vedras for allowing the study and sampling of Cova da Moura collection. We are grateful to the Orkney Museum for allowing access to samples from Orkney, and particularly to G. Drinkall for facilitating this work. We thank the Great North Museum: Hancock, the Society of Antiquaries of Newcastle upon Tyne, and Sunderland Museum for sharing samples. We are grateful to E. Willerslev for sharing several dozen samples that were analyzed in this study and for supporting several co-authors at the Centre for GeoGenetics at the University of Copenhagen who worked on this project. We are grateful for institutional support (grant RVO:67985912) from the Institute of Archaeology, Czech Academy of Sciences. G.K. was supported by Momentum Mobility Research Group of the Hungarian Academy of Sciences. This work was supported by the Wellcome Trust [100713/Z/12/Z]. D.F. was supported by an Irish Research Council grant GOIPG/2013/36. P.W.S., J.K. and A.M. were supported by the Heidelberg Academy of Sciences (WIN project “Times of Upheaval”). C.L.-F. was supported by a grant from FEDER and Ministry of Economy and Competitiveness (BFU2015-64699-P) of Spain. D.R. was supported by US National Science Foundation HOMINID grant BCS-1032255, US National Institutes of Health grant GM100233, and is an investigator of the Howard Hughes Medical Institute.

## Author Contributions

S.B., M.E.A, N.R., A.S.-N., A.M., N.B., M.F., E.H., M.M., J.O., K.S., R.P., J.K., W.H., I.B. and D.R. performed or supervised wet laboratory work. G.T.C. undertook the radiocarbon dating of a large fraction of the British samples. I.A., K.K., A.B., K.W.A., A.A.F., E.B., M.B.-B., D.B., C.B., C.Bo., L.B., T.A., L.Bü., S.C., L.C.N., O.E.C., G.C., B.C., A.D., K.E.D., N.D., M.E., C.E., M.K., J.F.F., H.F., C.F., M.G., R.G.P., M.H.-U., E.Had., G.H., N.J., T.K., K.M., S.P., P.L., O.L., A.L., J.L.M., T.M., J.I.M, K.Mc., M.B.G., A.Mo., G.K., V.K., A.C., R.Pa., A.E., K.Kö., T.H., J.L.C., C.L., M.P.P., P.W., T.D.P., P.P., P.-J.R., P.R., R.R., M.A.R.G., A.S., J.S., A.M.S., V.S., L.V., J.Z., D.C., T.Hi., V.H., A.Sh., K.-G.S., P.W.S., R.P., J.K., W.H., I.B., C.L.- F. and D.R. assembled archaeological material. I.O., S.M., T.B., A.M., E.A., M.L., I.L., N.P., Y.D., Z.F., D.F., P.d.K., M.G.T. and D.R. analysed or supervised analysis of data. I.O., C.L.-F. and D. R. wrote the manuscript with input from all co-authors.

## Supplementary Tables

**Supplementary Table 1**. Ancient individuals included in this study.

**Supplementary Table 2**. Mitochondrial haplogroup calls for individuals with newly reported data.

**Supplementary Table 3**. Y-chromosome calls for males with newly reported data.

---

**Extended Data Figure 1.**
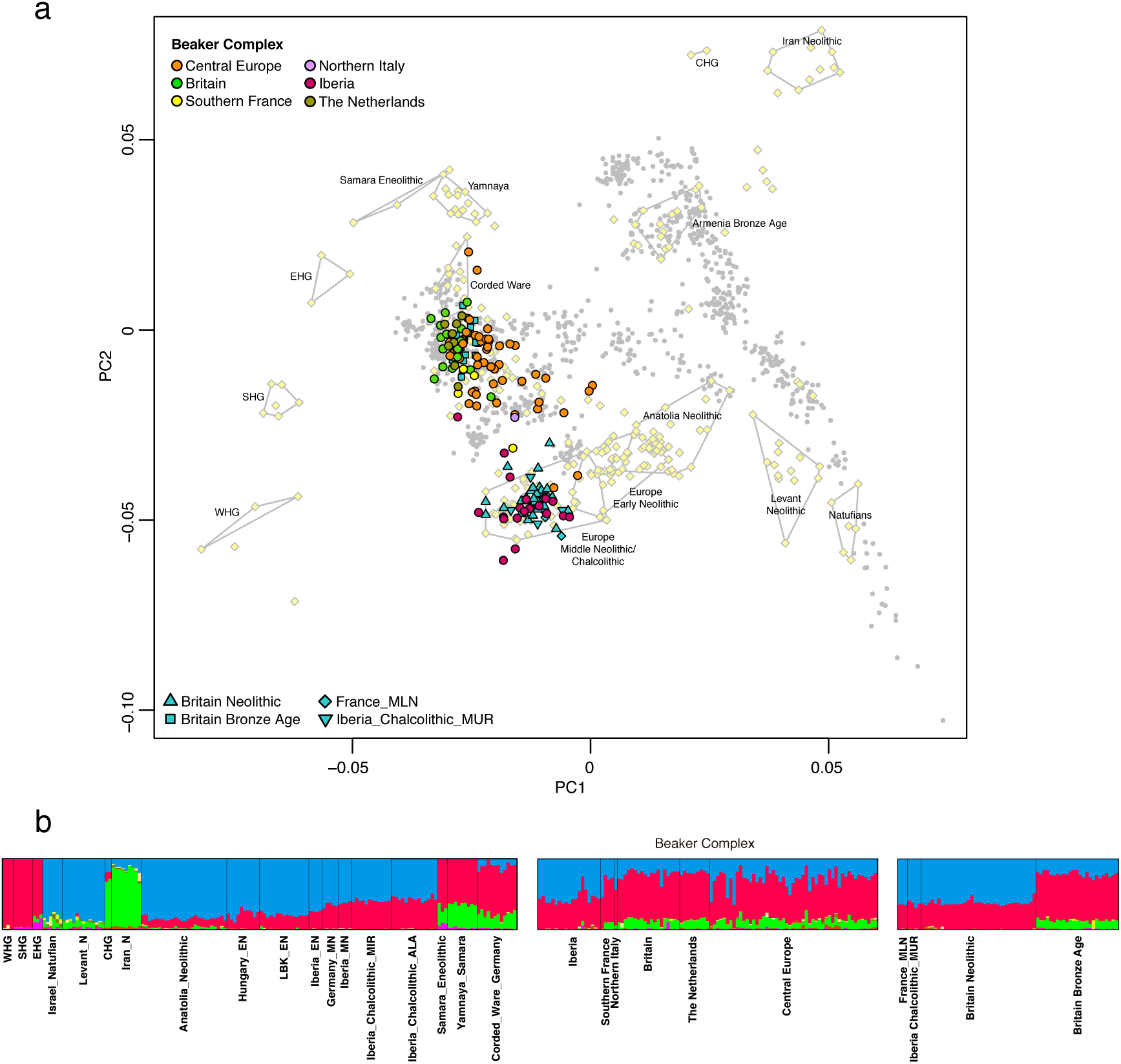
Population structure. **a,** Principal component analysis of 990 present-day West Eurasian individuals (grey dots), with previously published (pale yellow) and new ancient samples projected onto the first two principal components. **b**, ADMIXTURE clustering analysis with k=8 showing ancient individuals. E/M/MLN, Early/Middle/Middle Late Neolithic; W/E/S/CHG, Western/Eastern/Scandinavian/Caucasus hunter-gatherers.

**Extended Data Figure 2.**
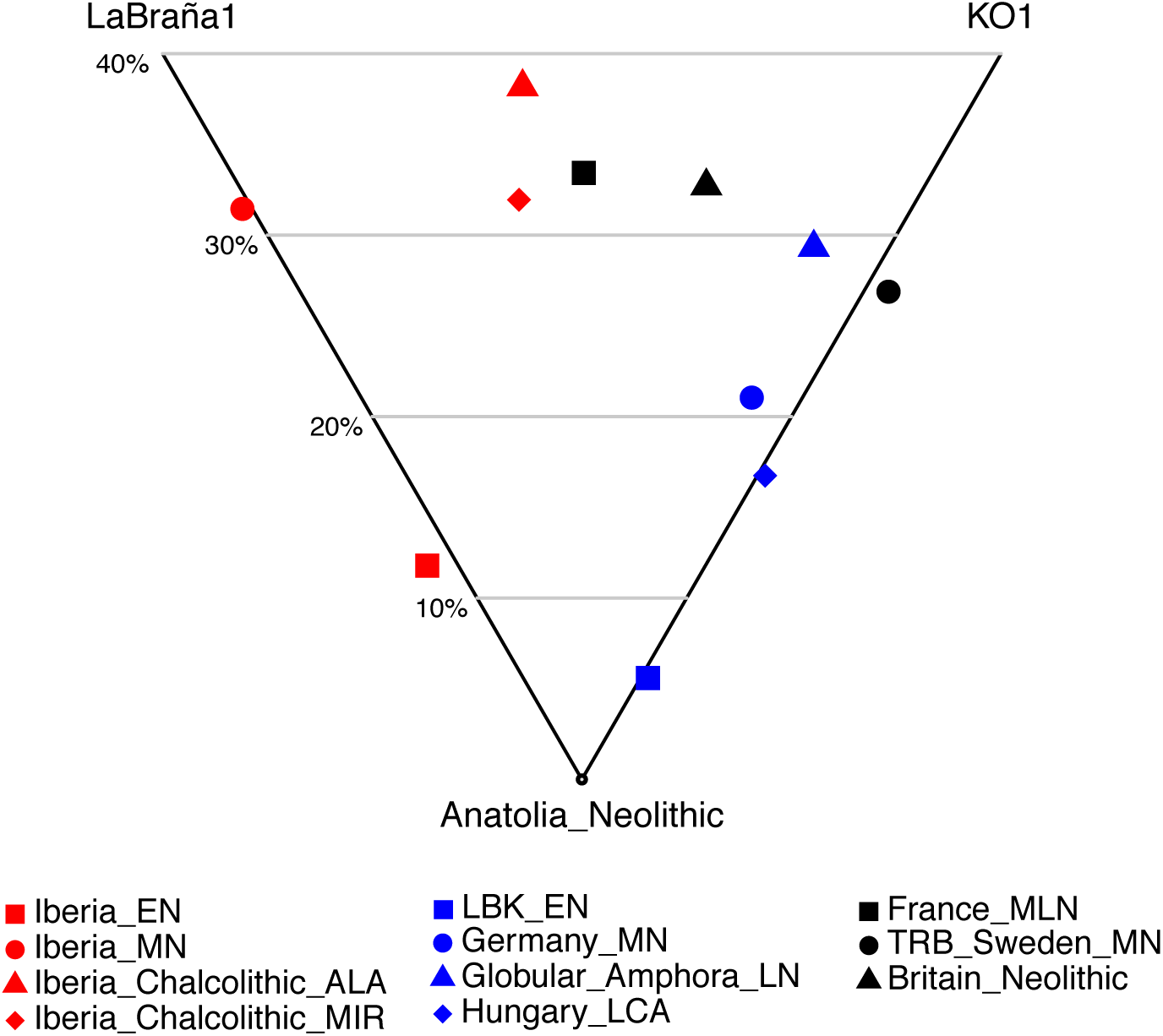
Hunter-gatherer affinities in Neolithic/Copper Age Europe. Differential affinity to hunter-gatherer individuals (LaBraña1^36^ from Spain and KO1^39^ from Hungary) in European populations before the emergence of the Beaker Complex. See Supplementary Information, section 6 for mixture proportions and standard errors computed with *qpAdm*.

**Extended Data Figure 3.**
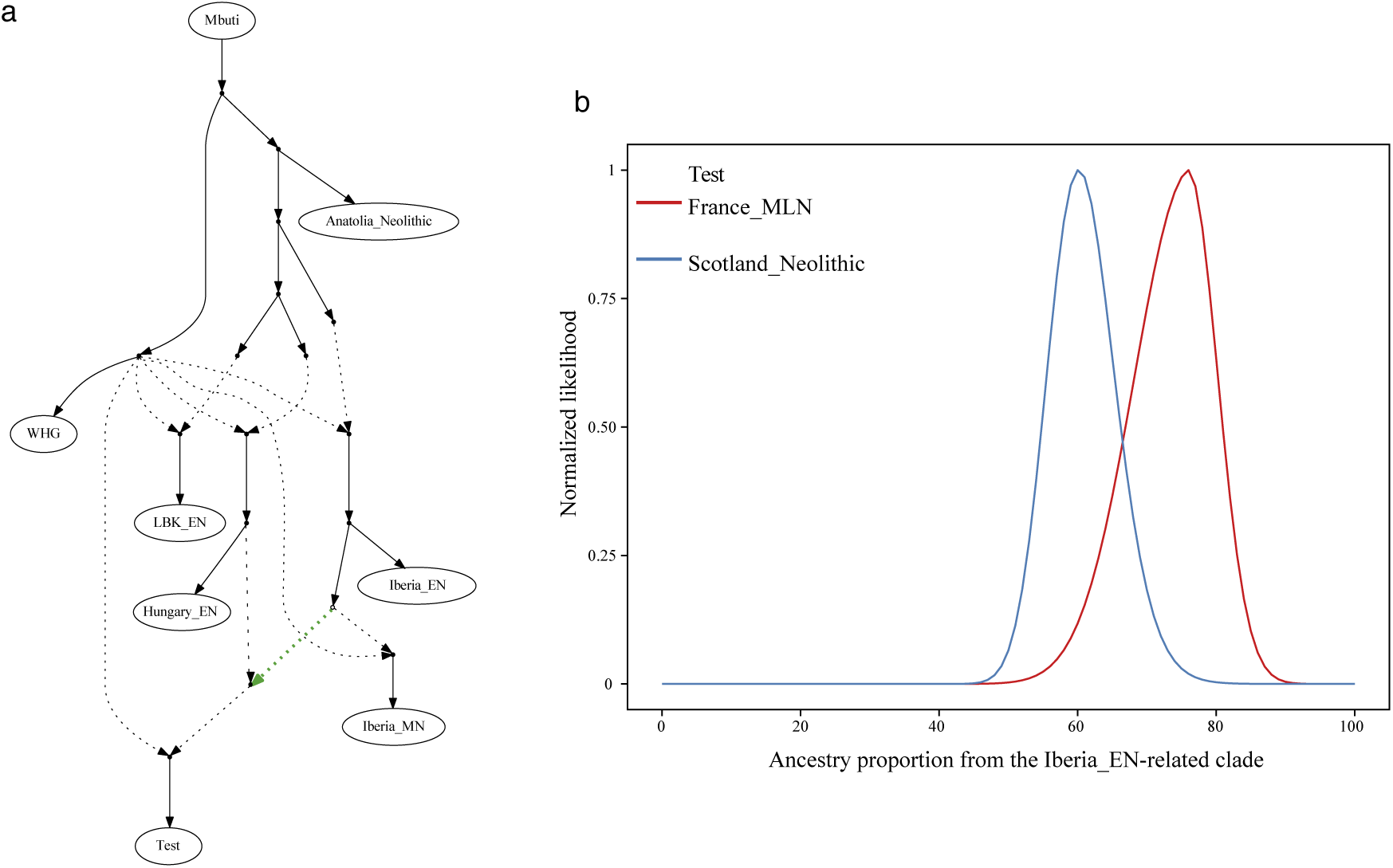
Modelling the relationships between Neolithic populations. **a,** Admixture graph fitting a *Test* population as a mixture of sources related to both Iberia_EN and Hungary_EN. **b,** Likelihood distribution for models with different proportions of the source related to Iberia_EN (green admixture edge in (a)) when *Test* is Great Britain_Neolithic or France_MLN.

**Extended Data Figure 4.**
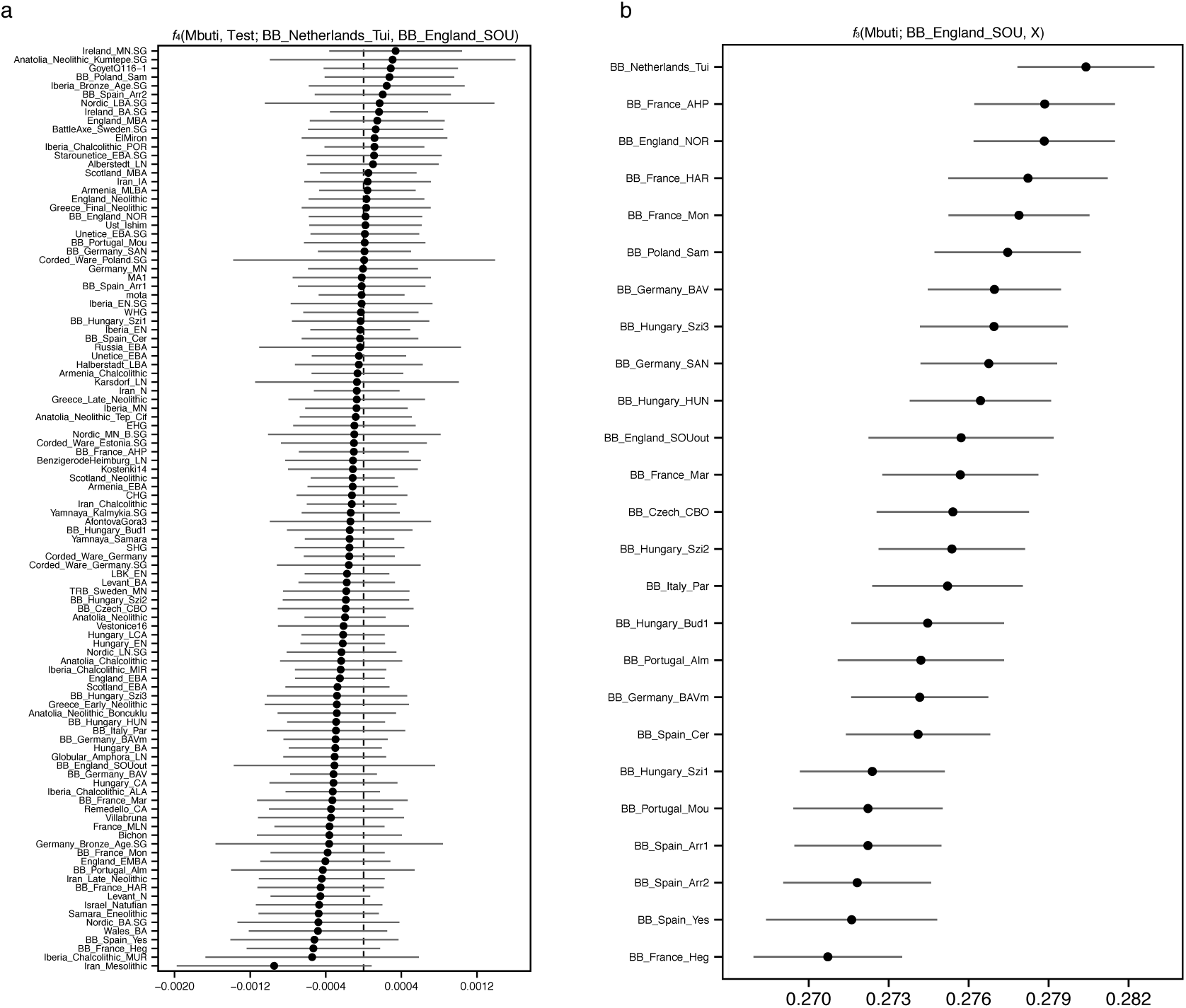
Genetic affinity between Beaker complex individuals from southern Great Britain and the Netherlands. **a**, *f-*statistics of the form *f*_4_(Mbuti, Test; BB_Netherlands_Tui, BB_Great Britain_SOU). Negative values indicate that Test is closer to BB_Netherlands_Tui than to BB_Great Britain_SOU, and the opposite for positive values. Error bars represent ±3 standard errors. **b**, Outgroup-*f*_3_ statistics of the form *f*_3_(Mbuti; BB_Great Britain_SOU, X) measuring shared genetic drift between BB_Great Britain_SOU and other Beaker Complex groups. Error bars represent ±1 standard errors.

**Extended Data Figure 5.**
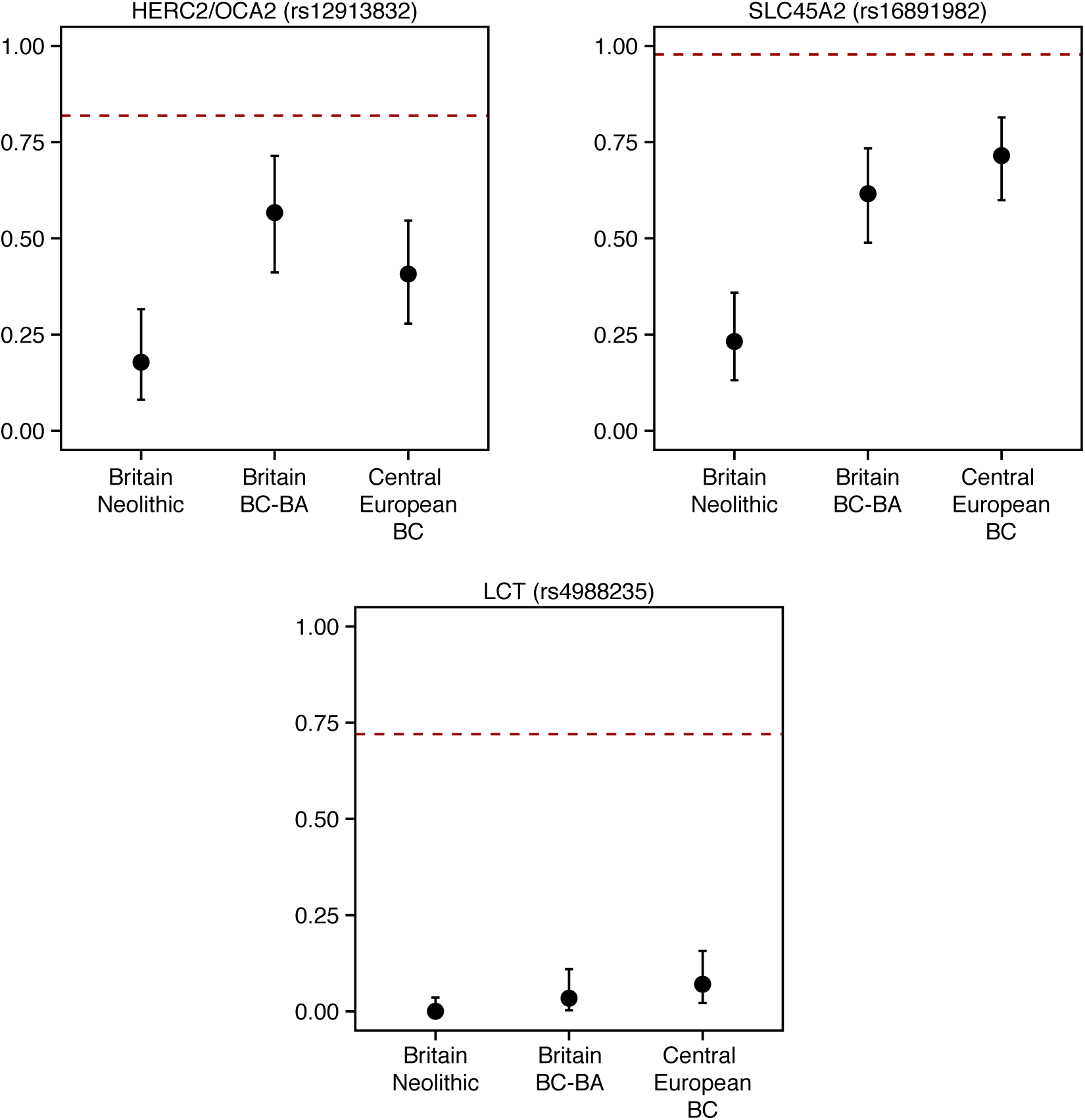
Derived allele frequencies at three SNPs of functional importance. Error bars represent 1.9-log-likelihood support interval. The red dashed lines show allele frequencies in the 1000 Genomes GBR population (present-day people from Great Britain). BC, Beaker Complex; BA, Bronze Age.

**Extended Data Table 1.** 62 Newly reported radiocarbon dates.

| Sample | Date | Location | Country |
| --- | --- | --- | --- |
| I4145 | 2279–2033 calBCE (3740±35 BP, Poz-84460) | Kněževes | Czech Republic |
| I1392 | 2832–2476 calBCE (4047±29 BP, MAMS-25935) | Hégenheim Necropole, Haut-Rhin | France |
| I4144 | 2572–2512 calBCE (3955±35 BP, Poz-84553) | Osterhofen-Altenmarkt | Germany |
| E09537_d | 2471–2300 calBCE (3909±29 BP, MAMS 29074) | Unterer Talweg 58-62, Augsburg, Bavaria | Germany |
| I4249 | 2336–2141 calBCE (3802±26 BP, BRAMS1217) | Irlbach LKR | Germany |
| E09538 | 2464–2212 calBCE (3870±30 BP, MAMS 29075) | Unterer Talweg 58-62, Augsburg, Bavaria | Germany |
| I3592 | 2458–2204 calBCE (3844±33 BP, BRAMS-1218) | Alburg-Lerchenhaid, Spedition Häring, Bavaria | Germany |
| I4250 | 2434–2150 calBCE (3825±26 BP, BRAMS1219) | Irlbach LKR | Germany |
| I3593 | 2398–2146 calBCE (3817±26 BP, BRAMS-1215) | Alburg-Lerchenhaid, Spedition Häring, Bavaria | Germany |
| I3590 | 2339–2143 calBCE (3802±26 BP, BRAMS-1217) | Alburg-Lerchenhaid, Spedition Häring, Bavaria | Germany |
| I2657 | 3952–3781 calBCE (5052±30 BP, SUERC-68701) | Macarthur Cave | Great Britain |
| I2633 | 3766–3642 calBCE (4911±32 BP, SUERC-68634) | Tulloch of Assery B | Great Britain |
| I2659 | 3762–3644 calBCE (4914±27 BP, SUERC-68702) | Distillery Cave | Great Britain |
| I2691 | 3701–3640 calBCE (4881±25 BP, SUERC-68704) | Distillery Cave | Great Britain |
| I2796 | 3706–3536 calBCE (4856±33 BP, SUERC-69074) | Point of Cott, Orkney | Great Britain |
| I2634 | 3704–3535 calBCE (4851±34 BP, SUERC-68638) | Tulach an t'Sionnach | Great Britain |
| I2635 | 3653–3390 calBCE (4796±37 BP, SUERC-68639) | Tulloch of Assery A | Great Britain |
| I2636 | 3520–3362 calBCE (4651±33 BP, SUERC-68640) | Holm of Papa Westray North | Great Britain |
| I2988 | 3517–3362 calBCE (4645±29 BP, SUERC-68711) | Clachaig | Great Britain |
| I2660 | 3514–3353 calBCE (4631±29 BP, SUERC-68703) | Distillery Cave | Great Britain |
| I2650 | 3500–3360 calBCE (4754±36 BP, SUERC-68642) | Holm of Papa Westray North | Great Britain |
| I2637 | 3510–3340 calBCE (4697±33 BP, SUERC-68641) | Holm of Papa Westray North | Great Britain |
| I2605 | 3632–3373 calBCE (4710±35 BP, Poz-83483) | Eton Rowing Course | Great Britain |
| I2980 | 3361–3102 calBCE (4530±33 BP, SUERC-69073) | Point of Cott, Orkney | Great Britain |
| I2651 | 3330–3090 calBCE (4525±36 BP, SUERC-68643) | Holm of Papa Westray North | Great Britain |
| I3085 | 3339–3027 calBCE (4471±29 BP, SUERC-68724) | Isbister, Orkney | Great Britain |
| I2978 | 3336–3024 calBCE (4464±29 BP, SUERC-68725) | Isbister, Orkney | Great Britain |
| I2934 | 3327–3036 calBCE (4466±33 BP, SUERC-69071) | Isbister, Orkney | Great Britain |
| I2935 | 3336–3012 calBCE (4451±29 BP, SUERC-68723) | Isbister, Orkney | Great Britain |
| I2979 | 3334–2942 calBCE (4447±29 BP, SUERC-68726) | Isbister, Orkney | Great Britain |
| I2631 | 3098–2907 calBCE (4384±36 BP, SUERC-68633) | Quoyness | Great Britain |
| I2933 | 3011–2886 calBCE (4309±29 BP, SUERC-68722) | Isbister, Orkney | Great Britain |
| I2977 | 3009–2764 calBCE (4275±33 BP, SUERC-69072) | Isbister, Orkney | Great Britain |
| I2630 | 2581–2464 calBCE (3999±32 BP, SUERC-68632) | Isbister, Orkney | Great Britain |
| I2932 | 2571–2348 calBCE (3962±29 BP, SUERC-68721) | Isbister, Orkney | Great Britain |
| I2612 | 2465–2209 calBCE (3865±35 BP, Poz-83492) | Hasting Hill, Sunderland, Tyne and Wear | Great Britain |
| I2416 | 2470–2285 calBC (3830±30 BP, Beta-432804) | Amesbury Down, Wiltshire | Great Britain |
| I2418 | 2440–2200 calBCE (3835±25 BP, NZA-32788) | Amesbury Down, Wiltshire | Great Britain |
| I2565 | 2470–2140 calBCE (3829±38 BP, OxA-13562) | Amesbury Down, Wiltshire | Great Britain |
| I2459 | 2460–2140 calBCE (3829±30 BP, SUERC-54823) | Amesbury Down, Wiltshire | Great Britain |
| I2457 | 2480–2280 calBCE (3890±30 BP, SUERC-36210) | Amesbury Down, Wiltshire | Great Britain |
| I2457 | 2200–2031 calBCE (3717±28 BP, SUERC-69975) | Amesbury Down, Wiltshire | Great Britain |
| I2453 | 2289–2041 calBCE (3760±35 BP, Poz-83404) | West Deeping | Great Britain |
| I2445 | 2137–1930 calBCE (3650±35 BP, Poz-83407) | Yarnton | Great Britain |
| I2596 | 2280–2030 calBCE (3739±30 BP, NZA-32484) | Amesbury Down, Wiltshire | Great Britain |
| I2566 | 2210–2030 calBCE (3734±25 BP, NZA-32490) | Amesbury Down, Wiltshire | Great Britain |
| I2452 | 2195–1920 calBCE (3700±30 BP, Beta-444979) | Dairy Farm, Willington | Great Britain |
| I2452 | 2277–2030 calBCE (3735±35 BP, Poz-83405) | Dairy Farm, Willington | Great Britain |
| I2598 | 2140–1940 calBCE (3664±30 BP, NZA-32494) | Amesbury Down, Wiltshire | Great Britain |
| I2460 | 2030–1820 calBCE (3575±27 BP, SUERC-53041) | Amesbury Down, Wiltshire | Great Britain |
| I2609 | 2023–1772 calBCE (3560±40 BP, Poz-83423) | Hexham Golf Course, Northumberland | Great Britain |
| I2610 | 1936–1746 calBCE (3515±35 BP, Poz-83498) | Summerhill, Blaydon, Tyne and Wear | Great Britain |
| I1775 | 1693–1600 calBCE (3344±27 BP, OxA-14308) | Great Orme Mines, Llandudno, North Wales | Great Britain |
| I2574 | 1415–1228 calBCE (3065±36 BP, SUERC-62072) | North Face Cave, Llandudno, North Wales | Great Britain |
| I2786 | 2459–2206 calBCE (3850±35 BP, Poz-83639) | Szigetszentmiklós, Felső Űrge-hegyi dűlő | Hungary |
| I2787 | 2458–2202 calBCE (3840±35 BP, Poz-83640) | Szigetszentmiklós, Felső Űrge-hegyi dűlő | Hungary |
| I2741 | 2458–2154 calBCE (3835±35 BP, Poz-83641) | Szigetszentmiklós, Felső Űrge-hegyi dűlő | Hungary |
| I4229 | 2289–2135 calBCE (3775±25 BP, PSU-1750) | Cova da Moura | Portugal |
| I0826 | 2833–2480 calBCE (4051±28 BP, MAMS-25940) | Paris Street, Cerdanyola, Barcelona | Spain |
| I0257 | 2571–2350 calBCE (3965±29 BP, MAMS-25937) | Paris Street, Cerdanyola, Barcelona | Spain |
| I0462 | 2566–2346 calBCE (3950±26 BP, MAMS-25936) | Arroyal I, Burgos | Spain |
| I0825 | 2474–2300 calBCE (3915±29 BP, MAMS-25939) | Paris Street, Cerdanyola, Barcelona | Spain |

**Extended Data Table 2.** Sites with new genome-wide data reported in this study.

| Site | N | Approx. date range (BCE) | Country |
| --- | --- | --- | --- |
| Eton Rowing Course | 1 | 3700–3300 | Great Britain |
| Banbury Lane | 3 | 3360–3100 | Great Britain |
| Totty Pot, Cheddar, Somerset | 1 | 2900–2400 | Great Britain |
| Abingdon Spring Road cemetery, Oxfordshire | 1 | 2500–2200 | Great Britain |
| Hasting Hill, Sunderland, Tyne and Wear | 2 | 2500–1700 | Great Britain |
| Amesbury Down, Wiltshire | 10 | 2500–1400 | Great Britain |
| Windmill Fields, Ingleby Barwick, County Durham | 2 | 2400–1900 | Great Britain |
| Yarnton | 4 | 2400–1900 | Great Britain |
| Staxton Beacon, Staxton, North Yorkshire | 1 | 2400–1600 | Great Britain |
| West Deeping | 1 | 2300–2000 | Great Britain |
| Dairy Farm, Willington, Bedfordshire | 1 | 2300–1900 | Great Britain |
| Over Narrows, Needingworth Quarry, Cambridgeshire | 2 | 2300–1900 | Great Britain |
| Porton Down, Wiltshire | 1 | 2200–1900 | Great Britain |
| Reaverhill, Barrasford, Northumberland | 1 | 2200–1900 | Great Britain |
| Trumpington Meadows | 2 | 2200–1900 | Great Britain |
| Hexham Golf Course, Northumberland | 1 | 2100–1700 | Great Britain |
| Summerhill, Blaydon, Tyne and Wear | 1 | 2000–1700 | Great Britain |
| Thanet, Kent | 3 | 2000–1600 | Great Britain |
| Boscombe Airfield, Wiltshire | 1 | 1800–1600 | Great Britain |
| Canada Farm, Sixpenny Handley, Dorset | 1 | 1500–1300 | Great Britain |
| Macarthur Cave | 1 | 4000–3700 | Great Britain |
| Distillery Cave | 3 | 3800–3300 | Great Britain |
| Raschoille Cave, Oban, Argyll and Bute | 6 | 3800–3200 | Great Britain |
| Tulach an t'Sionnach | 1 | 3700–3500 | Great Britain |
| Tulloch of Assery A | 1 | 3700–3300 | Great Britain |
| Point of Cott, Orkney | 2 | 3700–3100 | Great Britain |
| Clachaig | 1 | 3600–3300 | Great Britain |
| Holm of Papa Westray North | 4 | 3600–3000 | Great Britain |
| Isbister, Orkney | 10 | 3400–2300 | Great Britain |
| Quoyness | 1 | 3100–2900 | Great Britain |
| Dryburn Bridge | 2 | 2300–1800 | Great Britain |
| Eweford Cottages | 1 | 2200–1900 | Great Britain |
| Stenchme, Lop Ness, Orkney | 1 | 2000–1400 | Great Britain |
| Longniddry, Evergreen | 3 | 1500–1300 | Great Britain |
| Pabbay Mor | 1 | 1500–1200 | Great Britain |
| Great Orme Mines, Llandudno | 1 | 1700–1600 | Great Britain |
| North Face Cave, Llandudno | 1 | 1500–1200 | Great Britain |
| Kněževes | 2 | 2500–1900 | Czech Republic |
| Augsburg, Bavaria | 2 | 2800–1800 | Germany |
| Osterhofen-Altenmarkt, Bavaria | 1 | 2600–2000 | Germany |
| Unterer Talweg 58-62, Augsburg, Bavaria | 2 | 2500–2200 | Germany |
| Manching-Oberstimm, Bavaria | 1 | 2500–2000 | Germany |
| Irlbach, County of Straubing-Bogen, Bavaria | 2 | 2500–2000 | Germany |
| Bruck, City of Künzing, Bavaria | 2 | 2500–2000 | Germany |
| Hugo-Eckener-Straße, Augsburg | 3 | 2500–2000 | Germany |
| Unterer Talweg 85, Augsburg, Bavaria | 1 | 2400–2100 | Germany |
| Alburg, Lerchenhaid-Spedition Häring, Bavaria | 11 | 2300–2150 | Germany |
| Budakalász, Csajerszke (M0 Site 12) | 2 | 2500–2200 | Hungary |
| Szigetszentmiklós, Felső Űrge-hegyi dűlő | 4 | 2500–2100 | Hungary |
| Budapest-Békásmegyer | 2 | 2500–2000 | Hungary |
| Samborzec | 3 | 2600–2100 | Poland |
| De Tuithoorn, Oostwoud, Noord-Holland | 9 | 2300–1600 | The Netherlands |
| Via Guidorossi, Parma | 1 | 2200–1900 | Italy |
| Clos de Roque, Saint Maximin-la-Sainte-Baume | 3 | 4700–4400 | France |
| Collet Redon, La Couronne-Martigues | 1 | 3500–3100 | France |
| Hégenheim Necropole, Haut-Rhin | 1 | 2900–2400 | France |
| Dolmen of Villard, Lauzet-Ubaye | 2 | 2459–2242 | France |
| Sierentz, Les Villas d'Aurele, Haut-Rhin | 2 | 2600–2200 | France |
| La Fare, Forcalquier | 1 | 2500–2200 | France |
| Marlens, Sur les Barmes, Haute-Savoie | 1 | 2500–2100 | France |
| Mondelange, PAC de la Sente, Moselle | 2 | 2500–1900 | France |
| Rouffach, Haut-Rhin | 1 | 2400–2100 | France |
| Galeria da Cisterna, Almonda | 2 | 2500–2200 | Portugal |
| Cova da Moura | 1 | 2300–2100 | Portugal |
| Paris Street, Cerdanyola, Barcelona | 10 | 2900–2200 | Spain |
| Camino del Molino, Caravaca, Murcia | 4 | 2900–2100 | Spain |
| Camino de las Yeseras, San Fernando de Henares | 2 | 2280–1790 | Spain |
| Arroyal I, Burgos | 5 | 2600–2200 | Spain |

